# Integrating Bioinformatics and Machine Learning to Investigate the Mechanisms by Which Three Major Respiratory Infectious Diseases Exacerbate Heart Failure

**DOI:** 10.1101/2024.06.21.600018

**Authors:** Yiding Yu, Juan Zhang, Jingle Shi, Huajing Yuan, Quancheng Han, Yitao Xue, Yan Li

## Abstract

**Background:** Heart failure (HF) is a severe cardiovascular disease often worsened by respiratory infections like influenza, COVID-19, and community-acquired pneumonia (CAP). This study aims to uncover the molecular commonalities among these respiratory diseases and their impact on HF, identifying key mediating genes.

**Methods:** Datasets from the GEO database were analyzed for differential expression to find common molecular features of the three respiratory diseases. Weighted Gene Co-expression Network Analysis (WGCNA) was used to identify gene modules associated with HF. GO and KEGG enrichment analyses determined the biological processes and pathways involved in HF exacerbation by respiratory diseases. Key genes were screened using LASSO, RF, and SVM-RFE machine learning algorithms, with accuracy validated by ROC curves. Single-sample GSEA (ssGSEA) was performed, and the Drug Signature Database (DSigDB) was used for drug prediction. Immune infiltration analysis was conducted using CIBERSORT.

**Results:** We identified 51 characteristic genes of respiratory diseases and 10 potential genes exacerbating HF, primarily involved in innate immune response, inflammation, and coagulation pathways. Machine learning algorithms identified RSAD2 and IFI44L as key genes with high accuracy (AUC > 0.7). ssGSEA indicated RSAD2’s involvement in complement and coagulation cascades, while IFI44L is associated with myocardial contraction in HF progression. DSigDB predicted six potential therapeutic drugs. Immune infiltration analysis revealed significant differences in eight immune cell types between HF patients and healthy controls.

**Conclusion:** Our findings enhance the understanding of molecular interactions between respiratory diseases and heart failure, paving the way for future research and therapeutic strategies.

## INTRODUCTION

Heart failure is a global public health issue, affecting approximately 6.2 million adults in the United States alone, with its prevalence continuing to rise as the population ages [1]. When patients with heart failure experience respiratory infections, their rates of hospitalization and mortality significantly increase. Studies have shown that hospitalizations due to respiratory infectious diseases account for 15.3% of heart failure cases, and these patients face a 60% higher risk of in-hospital mortality [2]. Since the outbreak of the COVID-19 pandemic at the end of 2019, public health systems have been under immense pressure [3]. Although the prevalence of influenza decreased during the COVID-19 pandemic, it still imposed an additional burden on public health systems [4]. Furthermore, community-acquired pneumonia (CAP), one of the most common respiratory infections, has become more complex to treat during the COVID-19 pandemic due to co-infection with the SARS-CoV-2 virus [5]. Common symptoms among these respiratory disease patients, such as cough, dyspnea, fever, as well as inflammatory responses, T-cell exhaustion and dysfunction, and immune evasion mechanisms, can all exacerbate the cardiac burden in heart failure patients [7-8].

Research indicates that respiratory infectious diseases such as COVID-19, influenza, and CAP exacerbate heart failure, leading to increased morbidity and mortality [9]. For instance, COVID-19 is associated with a higher incidence of acute cardiac injury and heart failure, particularly in patients with pre-existing cardiovascular conditions [10]. Similarly, influenza and CAP can trigger acute decompensated heart failure, further complicating the clinical management of these patients [11]. These respiratory infectious diseases exacerbate the progression of heart failure through various mechanisms, including inducing systemic inflammatory responses, increasing the metabolic demands on the heart, and affecting cardiac structure and function. However, the specific interaction mechanisms between these diseases and heart failure remain unclear. Understanding the molecular mechanisms underlying these interactions is crucial for developing targeted therapies that can mitigate the impact of respiratory infections on heart failure.

In this study, we first identified specific genes associated with COVID-19, influenza, and CAP through differential expression analysis. Subsequently, using Weighted Gene Co-expression Network Analysis (WGCNA), we screened for module genes related to heart failure, thereby identifying genes that may play critical roles in the exacerbation of heart failure by respiratory infectious diseases. Through functional enrichment analysis, we further explored the roles of these genes in biological processes. To accurately identify key genes associated with the exacerbation of heart failure by respiratory infectious diseases, we employed three machine learning algorithms: Least Absolute Shrinkage and Selection Operator (LASSO), Random Forest (RF), and Support Vector Machine-Recursive Feature Elimination (SVM-RFE) for further screening of key genes. To assess the accuracy of our screening results, we validated them using external datasets and plotted ROC curves. We also performed single-gene Gene Set Enrichment Analysis (GSEA) for the key genes. Additionally, we conducted drug prediction based on the DSigDB database to identify potential therapeutic strategies aimed at alleviating the symptoms and cardiac burden of heart failure patients affected by respiratory infections. Finally, through immune infiltration analysis, we investigated the impact of immune responses induced by respiratory infectious diseases on myocardial failure, aiming to uncover the complex interaction mechanisms between these diseases. Figure 1 depicts the study flowchart.

**Figure 1:**
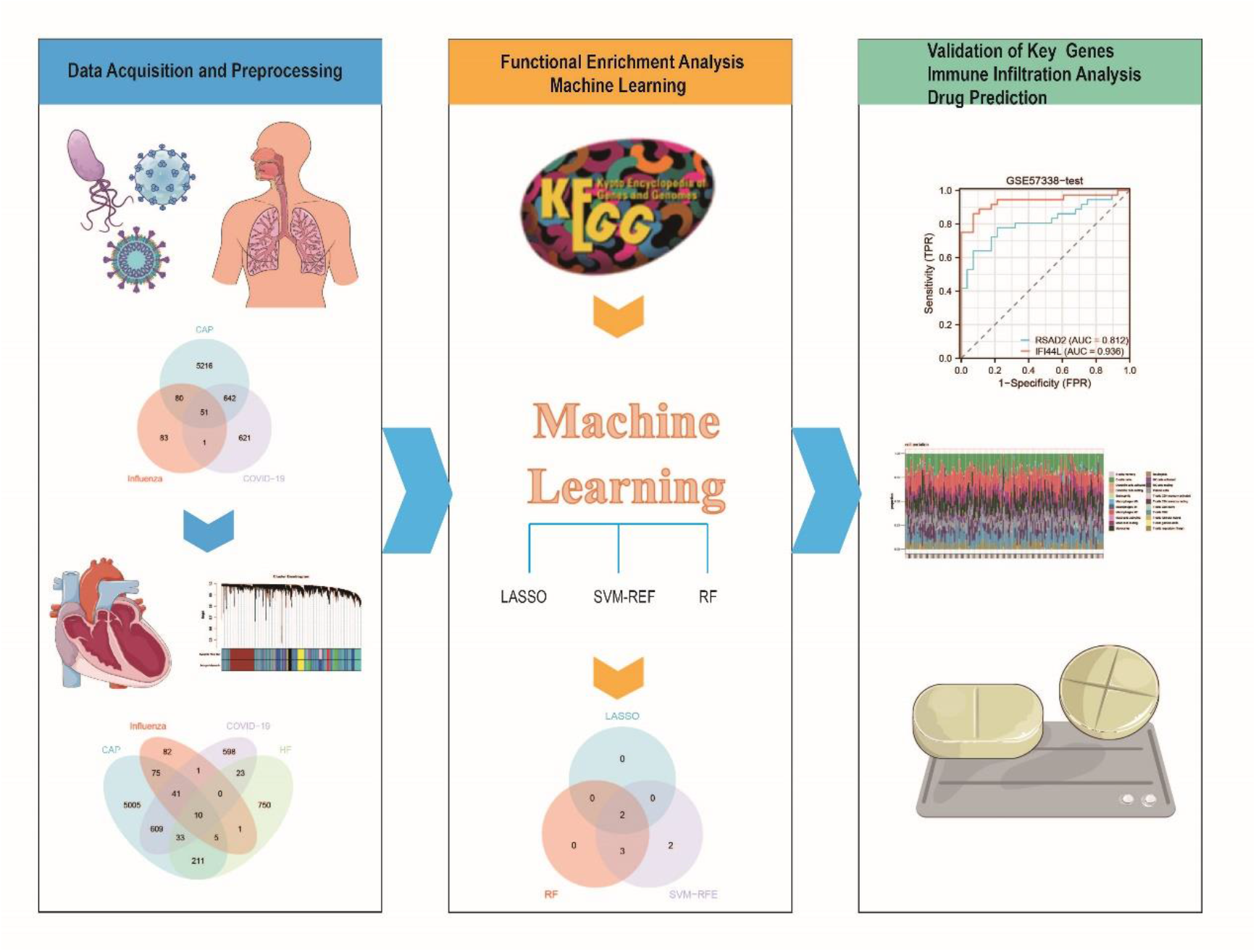
The study flowchart.

## MATERIALS AND METHODS

### Microarray Data

The data for this study were sourced from the publicly available GEO database. The datasets had obtained consent and ethical approval from the relevant participants, thus institutional review board approval was not required for this study [12]. For the heart failure dataset, we selected GSE57338, which includes left ventricular myocardial samples from 95 ischemic heart failure patients and 136 normal individuals [13]. We randomly selected 70% of the samples as the training set, with the remaining 30% as the test set. For external validation, we chose GSE5406, which comprises left ventricular myocardial samples from 108 ischemic heart failure patients and 16 normal individuals [14]. For the influenza dataset, we selected GSE111368, which includes whole blood samples from 229 influenza patients and 130 normal individuals [15]. For the COVID-19 dataset, we selected GSE157103, which includes leukocyte samples from 100 COVID-19 patients and 26 normal individuals [16]. For the community-acquired pneumonia dataset, we selected GSE196399, which includes leukocyte samples from 56 community-acquired pneumonia patients and 21 normal individuals [17]. For the validation of respiratory disease datasets, we selected GSE164805, GSE185576, and GSE94916 [18]. The dataset information is summarized in Table 1.

**Table 1:**
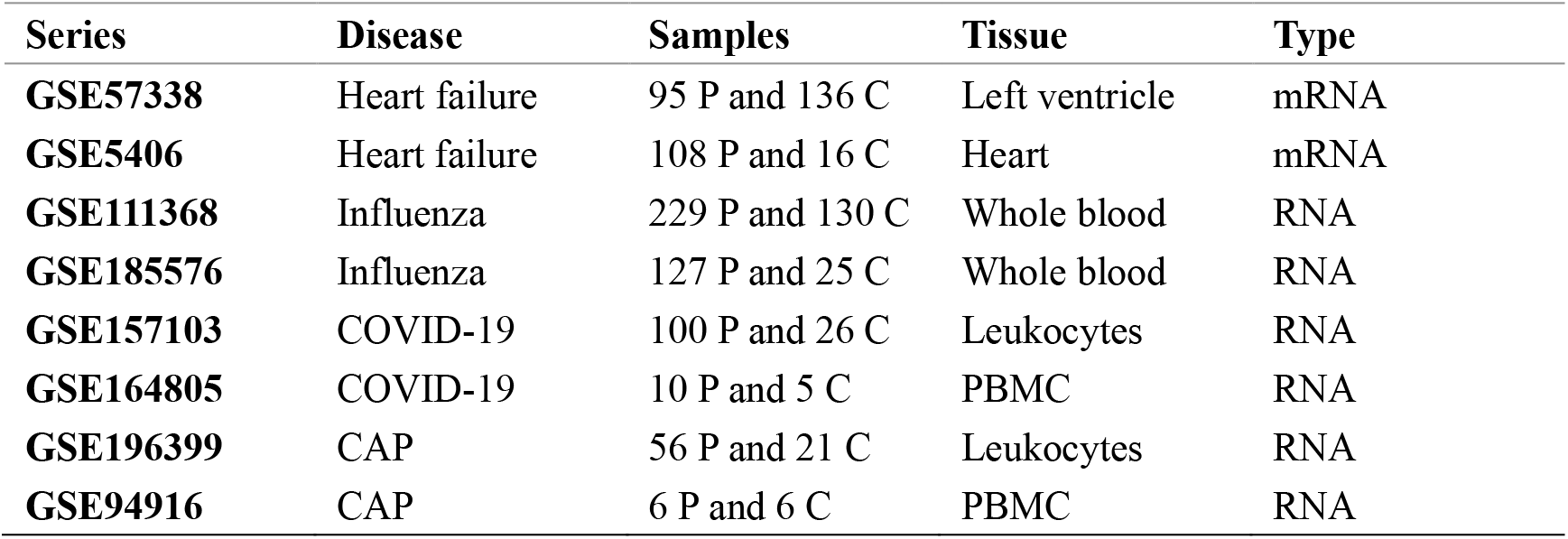
The details of disease datasets. P: Patient samples. C: Control group samples. CAP: Community-acquired pneumonia. PBMC: Peripheral blood mononuclear cells.

| Series | Disease | Samples | Tissue | Type |
| --- | --- | --- | --- | --- |
| **GSE57338** | Heart failure | 95 P and 136 C | Left ventricle | mRNA |
| **GSE5406** | Heart failure | 108 P and 16 C | Heart | mRNA |
| **GSE111368** | Influenza | 229 P and 130 C | Whole blood | RNA |
| **GSE185576** | Influenza | 127 P and 25 C | Whole blood | RNA |
| **GSE157103** | COVID-19 | 100 P and 26 C | Leukocytes | RNA |
| **GSE164805** | COVID-19 | 10 P and 5 C | PBMC | RNA |
| **GSE196399** | CAP | 56 P and 21 C | Leukocytes | RNA |
| **GSE94916** | CAP | 6 P and 6 C | PBMC | RNA |

### Data Processing and Differentially Expressed Gene Screening

We utilized R software (version 4.4.0) for data preprocessing. Probes corresponding to multiple molecules were removed. For probes corresponding to the same molecule, only the probe with the highest signal value was retained. We eliminated batch effects from the data and converted probe IDs to gene symbols based on the platform’s annotation files. Differential expression analysis for the respiratory disease datasets was performed using the limma package, selecting genes with p-values < 0.05 and |log2(FC)| ≥ 1 as differentially expressed genes [19].

Weighted Gene Co-Expression Network Analysis and Module Gene Selection We employed Weighted Gene Co-Expression Network Analysis (WGCNA) to explore gene modules associated with heart failure [20]. Using a threshold of 0.5, we filtered out unqualified genes and samples via the goodSamplesGenes function, and then constructed a scale-free co-expression network. Subsequently, we set the soft-thresholding power to β = 30 and scale-free R2 = 0.9 to calculate adjacency, converting adjacency into a Topological Overlap Matrix (TOM) to determine gene ratios and dissimilarities. Genes with similar expression profiles were grouped into gene modules using average linkage hierarchical clustering. We preferred larger modules and thus set the minimum module size to 200. Finally, we calculated the dissimilarity of module eigengenes, selected the cut-off line of the module dendrogram to combine several modules for further study, and visualized the eigengene network.

### Functional Enrichment Analysis

To understand the common molecular characteristics of the three respiratory infectious diseases (COVID-19, influenza, and CAP) and their potential impact on heart failure, we first intersected the differentially expressed genes of the three respiratory infectious diseases to identify their shared genes. Subsequently, we intersected these shared genes with the key modules identified by WGCNA for heart failure, thereby screening for candidate genes that may play a role in the exacerbation of heart failure by respiratory infectious diseases. To explore the biological processes and functions involving these genes, we performed Gene Ontology (GO) and Kyoto Encyclopedia of Genes and Genomes (KEGG) enrichment analyses using the clusterProfiler package [21-22].

### Machine Learning

We employed three machine learning algorithms—LASSO, RF, and SVM-RFE—to further screen for key genes involved in the exacerbation of heart failure by respiratory infectious diseases [23-25]. The LASSO algorithm was executed using the glmnet package, with ten-fold cross-validation to select prominent genes. The RF algorithm was performed using the randomForest package, selecting genes with higher scores as candidate genes. The SVM-RFE algorithm was implemented using the e1071 package, choosing the number of genes with the highest accuracy as candidate genes. After computation, we selected the intersection of the three sets as the key genes.

### Key Genes Verification

To ensure the accuracy and reliability of the identified key genes, we validated them in the heart failure and respiratory disease validation datasets. We constructed Receiver Operating Characteristic (ROC) curves to evaluate the diagnostic accuracy of these key genes. The value was quantified by calculating the Area Under the Curve (AUC) of the ROC, with AUC > 0.7 considered accurate and reliable.

### Single Sample Gene Set Enrichment Analysis

We performed single-gene Gene Set Enrichment Analysis (GSEA) for the key genes using the clusterProfiler package to explore the functions involved in the exacerbation of heart failure by respiratory infectious diseases [26].

### Drug Prediction

The Drug Signatures Database (DSigDB) is a comprehensive database designed to facilitate the association studies between gene sets and drug characteristics [27]. It includes the effects of various drugs on cells and the gene expression changes induced by these drugs, providing valuable resources for drug repurposing and new drug discovery. We compiled a list of the previously identified key genes and utilized the DSigDB database to predict potential drug molecules that may alleviate symptoms and cardiac burden in heart failure patients, particularly those exacerbated by respiratory infections.

### Immune Infiltration Analysis

We utilized the CIBERSORT package to assess the content of immune cells and stromal cells in myocardial samples, aiming to depict the cellular heterogeneity landscape of myocardial expression profiles and complete the immune cell infiltration analysis [28]. Bar plots were used to visualize the proportion of each type of immune cell in different samples. The Wilcoxon rank-sum test was employed to compare the differences in cell distribution between the heart failure and normal groups, with a cut-off value set at p < 0.05.

## RESULTS

### Identification of Differentially Expressed Genes

After completing data processing and differential expression analysis, we identified 215 differentially expressed genes for influenza, 1315 for COVID-19, and 5989 for community-acquired pneumonia. By intersecting the differentially expressed genes of the three diseases, we obtained 51 specific genes associated with respiratory infectious diseases, as illustrated in Figure 2A.

**Figure 2:**
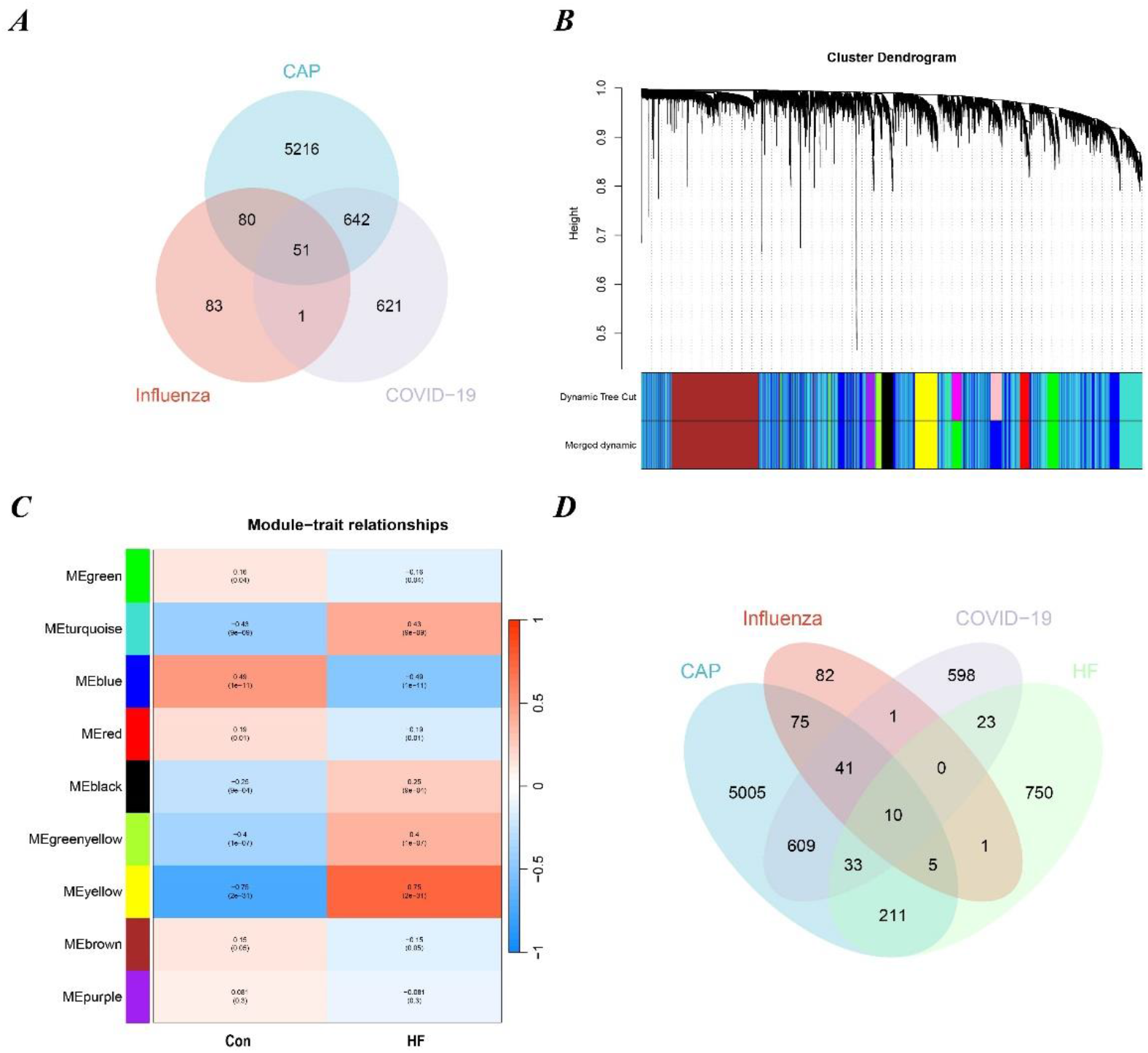
Results of Differentially Expressed Genes and WGCNA. (A) 51 specific genes associated with respiratory infectious diseases. (B) Gene co-expression modules represented by different colors under the gene tree. (C) 9 gene co-expression modules. (D) Intersection of genes in respiratory infectious diseases and heart failure.

### Weighted Gene Co-Expression Network Analysis and Key Module Identification

We employed WGCNA to identify the most relevant modules in heart failure. Ultimately, we identified nine gene co-expression modules. Among them, the yellow module exhibited the highest correlation with HF (correlation coefficient = 0.75, p = 2e-31), comprising a total of 1033 genes. Therefore, we selected the yellow module as the key module for subsequent analysis. By intersecting the module genes with the respiratory disease-specific genes, we identified 10 genes that may play critical roles in the exacerbation of heart failure by respiratory infectious diseases. The relevant results are illustrated in Figures 2B-2D.

### Functional Enrichment Analysis

We performed GO and KEGG enrichment analyses on the 51 respiratory disease-specific genes and the 10 genes that may play critical roles in the exacerbation of heart failure by respiratory infectious diseases. The GO enrichment analysis categories included Biological Process (BP), Cellular Component (CC), and Molecular Function (MF).

For the 51 respiratory disease-specific genes, the BP terms were primarily associated with defense response to Gram-negative bacterium, antimicrobial humoral response, defense response to bacterium, antibacterial humoral response, and innate immune response in mucosa. The CC terms were mainly related to primary lysosome, azurophil granule, secretory granule lumen, cytoplasmic vesicle lumen, and vesicle lumen. The MF terms were predominantly linked to lipopolysaccharide binding, serine-type endopeptidase activity, serine-type peptidase activity, serine hydrolase activity, and heparin binding. KEGG enrichment analysis indicated that these 51 genes were mainly involved in Staphylococcus aureus infection, NOD-like receptor signaling pathway, Transcriptional misregulation in cancer, Neutrophil extracellular trap formation, and Hepatitis C.

For the 10 genes that may play critical roles in the exacerbation of heart failure by respiratory infectious diseases, the BP terms were primarily associated with response to virus, negative regulation of viral genome replication, regulation of viral genome replication, negative regulation of viral process, and defense response to virus. No significant results were obtained for CC terms. The MF terms were mainly linked to double-stranded RNA binding, caspase binding, immunoglobulin binding, GTP binding, and adenylyltransferase activity. KEGG enrichment analysis indicated that these genes were mainly involved in Hepatitis C, Influenza A, Measles, Coronavirus disease - COVID-19, and Epstein-Barr virus infection. The visualization results are shown in Figure 3.

**Figure 3:**
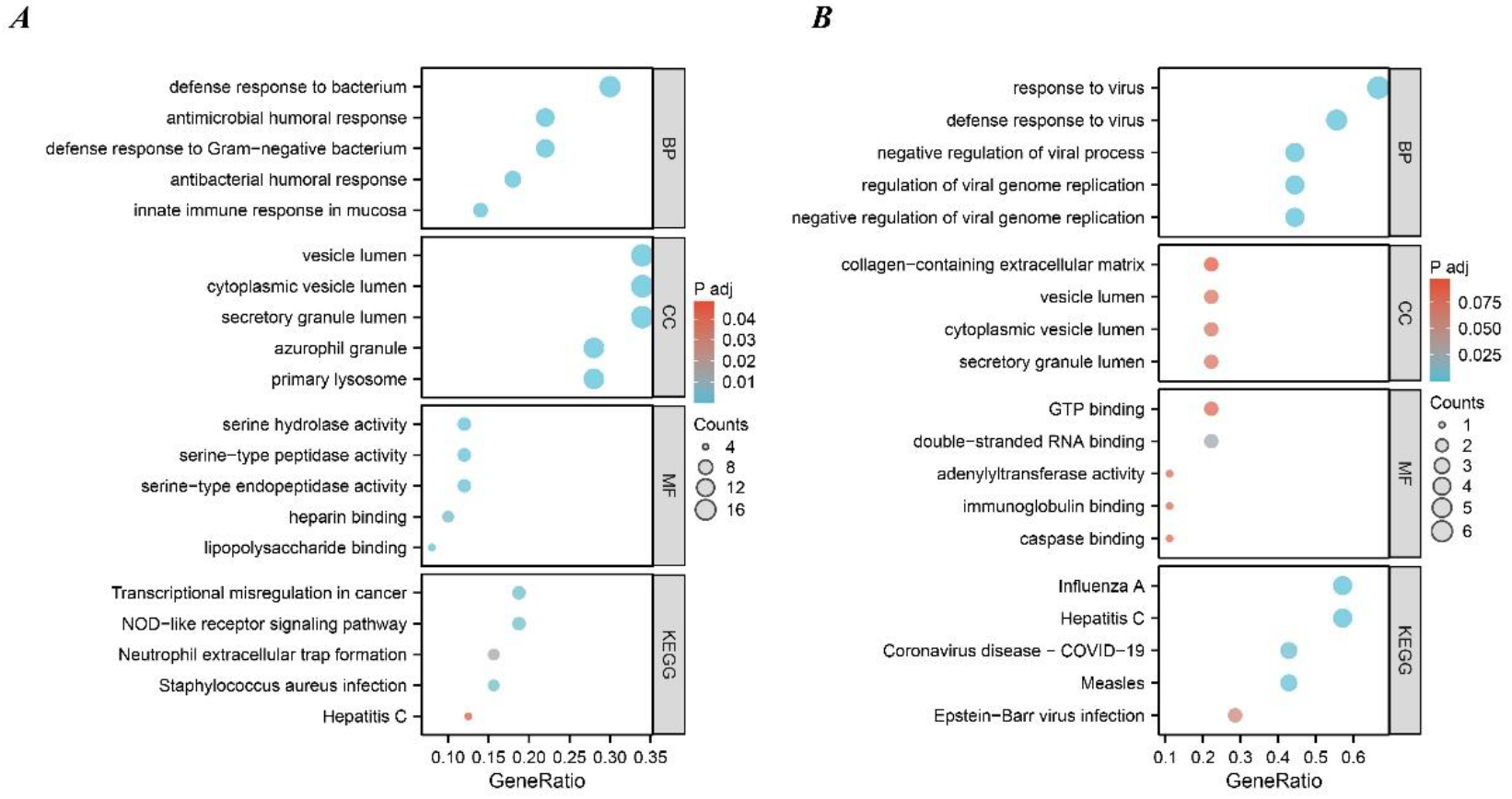
Functional Enrichment Analysis. (A) Enrichment analysis results of 51 specific genes associated with respiratory infectious diseases. (B) Enrichment analysis results of 10 key genes.

### Identification of Hub Genes via Machine Learning

We employed three machine learning algorithms—LASSO, RF, and SVM-RFE—to further screen for key genes. The LASSO algorithm identified 2 candidate genes. The RF algorithm ranked genes based on their importance, selecting genes with importance scores greater than 7 as candidate genes, resulting in 5 candidate genes. The SVM-RFE algorithm indicated that the highest accuracy and lowest error rate were achieved with 7 genes. Therefore, we selected the top 7 genes from the SVM-RFE algorithm as candidate genes. By intersecting the results of the three algorithms, we identified RSAD2 and IFI44L as the key genes. The visualization results are shown in Figure 4.

**Figure 4:**
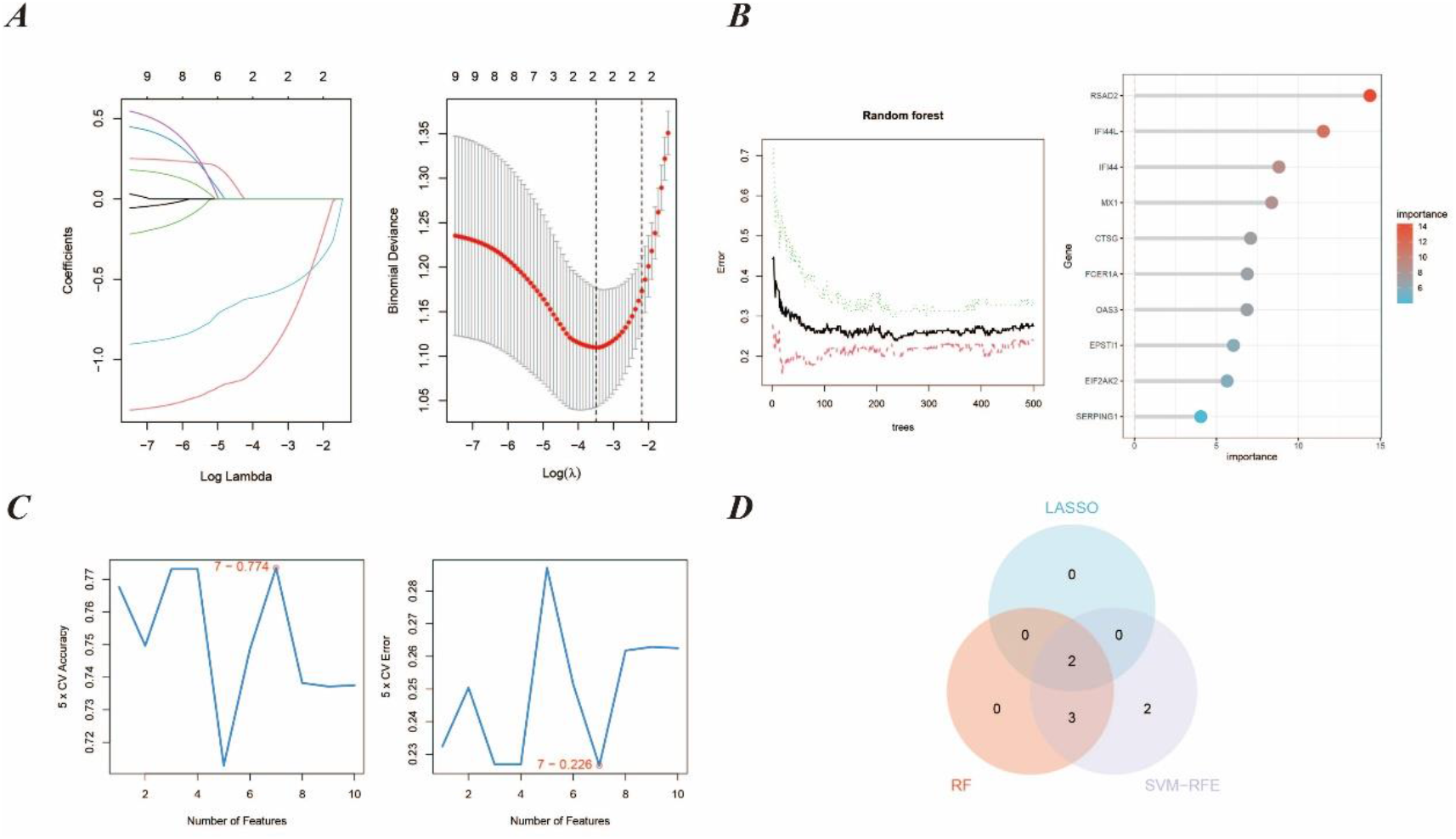
Machine learning in screening key genes. (A) Key genes screening in the Lasso model. (B) Key genes in the RF model. (C) Key genes in the SVM-RFE model. (D) Venn diagram shows that 2 key genes are identified via the above three algorithms.

### Key Genes Verification

We first verified the accuracy of RSAD2 and IFI44L in myocardial samples, including the internal validation set GSE57338 and the external validation set GSE5406. The results indicated that the AUCs of both genes were greater than 0.7 in both the internal and external validation sets. Subsequently, we validated these two genes in external datasets for influenza, COVID-19, and community-acquired pneumonia. The results showed that RSAD2 and IFI44L had high accuracy in the COVID-19 validation set, with AUCs greater than 0.7. In the influenza validation set, only RSAD2 had an AUC greater than 0.7. In the community-acquired pneumonia validation set, only IFI44L had an AUC greater than 0.7. The visualization results are shown in Figure 5.

**Figure 5:**
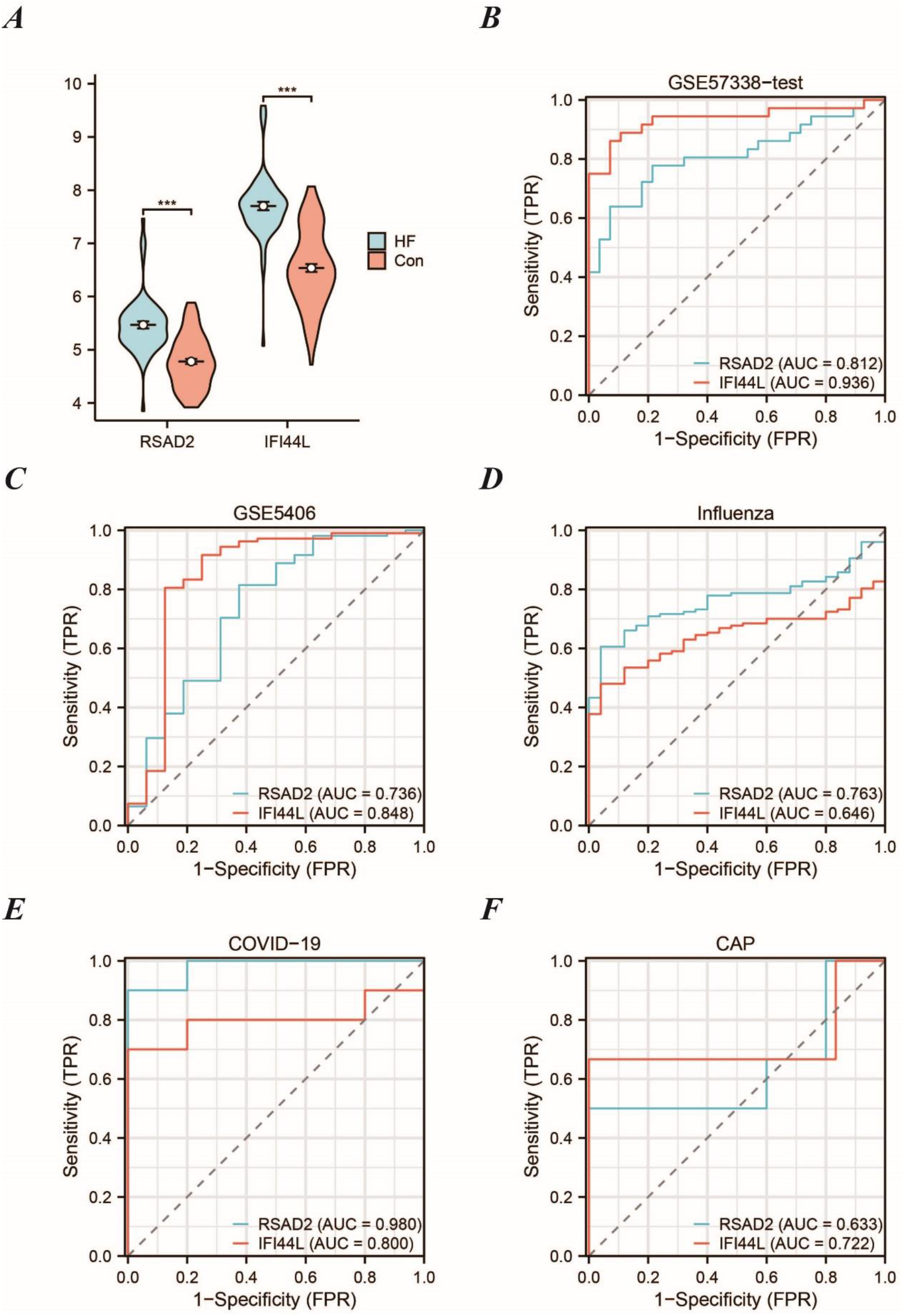
Verification of the accuracy of key genes. (A) Expression of key genes in HF patients relative to normal controls in dataset GSE57338-train. (B) The ROC curve of each key genes in GSE57338-test. (C) The ROC curve of each key genes in GSE5406. (D) The ROC curve of each key genes in influenza dataset. (E) The ROC curve of each key genes in COVID-19 dataset. (F) The ROC curve of each key genes in CAP dataset.

### Single Sample Gene Set Enrichment Analysis

We performed ssGSEA on RSAD2 and IFI44L. The results indicated that during the progression of heart failure (HF), RSAD2 is primarily involved in functions such as complement and coagulation cascades, interactions between viral proteins and cytokines and cytokine receptors, proteasome, and compound metabolism. IFI44L is mainly involved in myocardial contraction, the intestinal immune network for IgA production, and various compound metabolism functions. Detailed results are shown in Figure 6.

**Figure 6:**
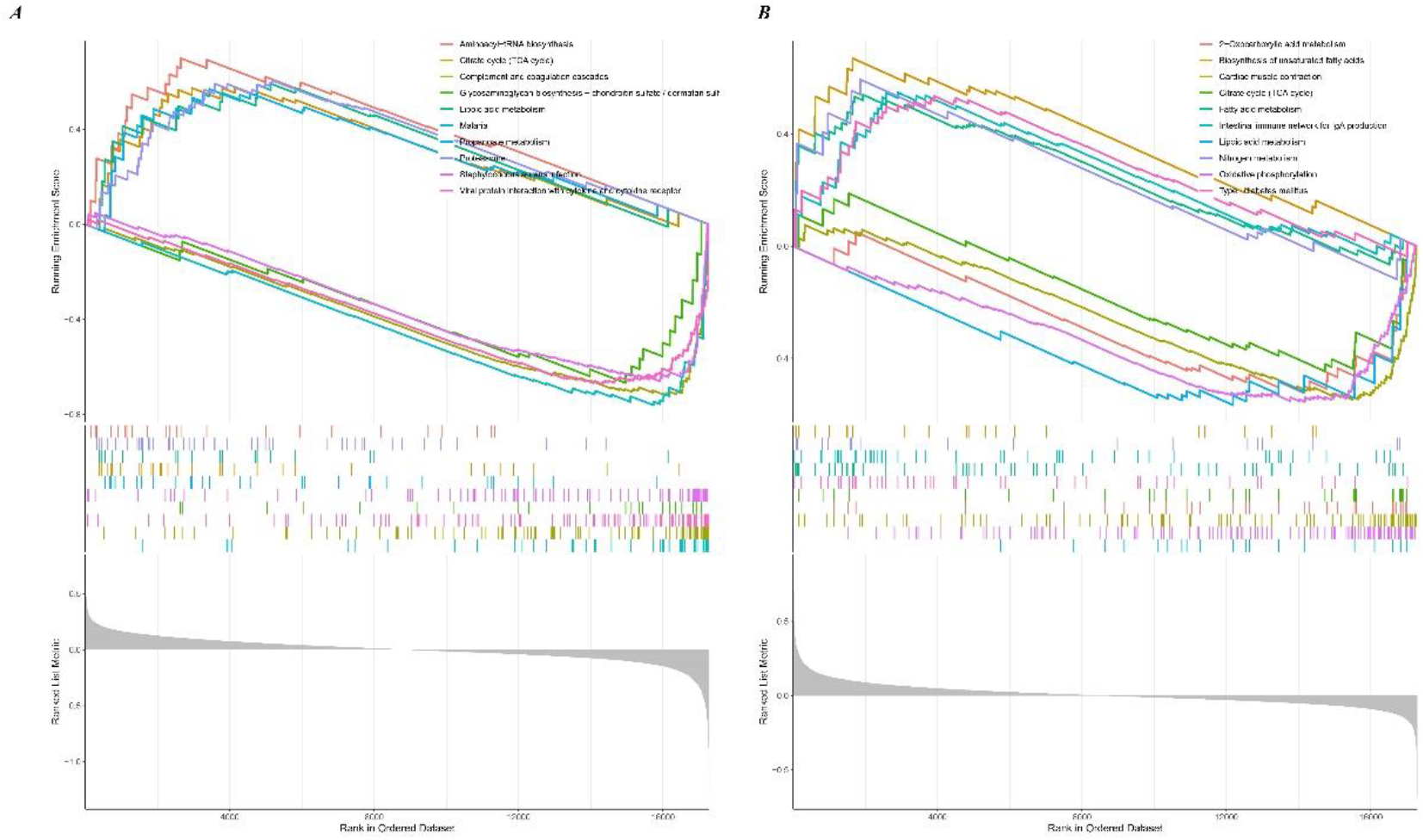
The results of ssGSEA Enrichment Analysis. (A) The results of RSAD2. (B) The results of IFI44L.

### Drug Prediction of Key Genes

We utilized the DSigDB database to predict potential drug molecules that may alleviate symptoms and cardiac burden in heart failure patients, particularly those exacerbated by respiratory infections. We selected drugs with an adjusted P-value less than 0.01, identifying a total of 6 drugs: acetohexamide, Gadodiamide hydrate, suloctidil, 3’-Azido-3’-deoxythymidine, testosterone enanthate, and tamoxifen. Detailed information is provided in Table 2.

**Table 2:** Details of predicted drugs.

| Term | Adjusted P-value | Combined Score | Genes |
| --- | --- | --- | --- |
| acetohexamide | 4.30E-05 | 542594.4 | RSAD2, IFI44L |
| Gadodiamide hydrate | 7.68E-05 | 491266.6 | RSAD2, IFI44L |
| suloctidil | 5.59E-04 | 393866.3 | RSAD2, IFI44L |
| 3'-Azido-3'-deoxythymidine | 0.002964 | 312489.9 | RSAD2, IFI44L |
| testosterone enanthate | 0.004821 | 282341 | RSAD2, IFI44L |
| tamoxifen | 0.009101 | 247038.9 | RSAD2, IFI44L |

### Immune Infiltration Analysis

We performed immune infiltration analysis on myocardial samples from the training set using the CIBERSORT algorithm. The bar plots clearly displayed the content of different subpopulations in each sample. We assessed the cellular composition heterogeneity between heart failure samples and healthy samples. The results indicated that there were significant differences in the infiltration of 8 types of immune cells. This may provide new insights into understanding the mechanisms by which respiratory infections exacerbate heart failure and could also offer potential regulatory points for the treatment of heart failure patients affected by respiratory infections. The visualization results are shown in Figure 7.

**Figure 7:**
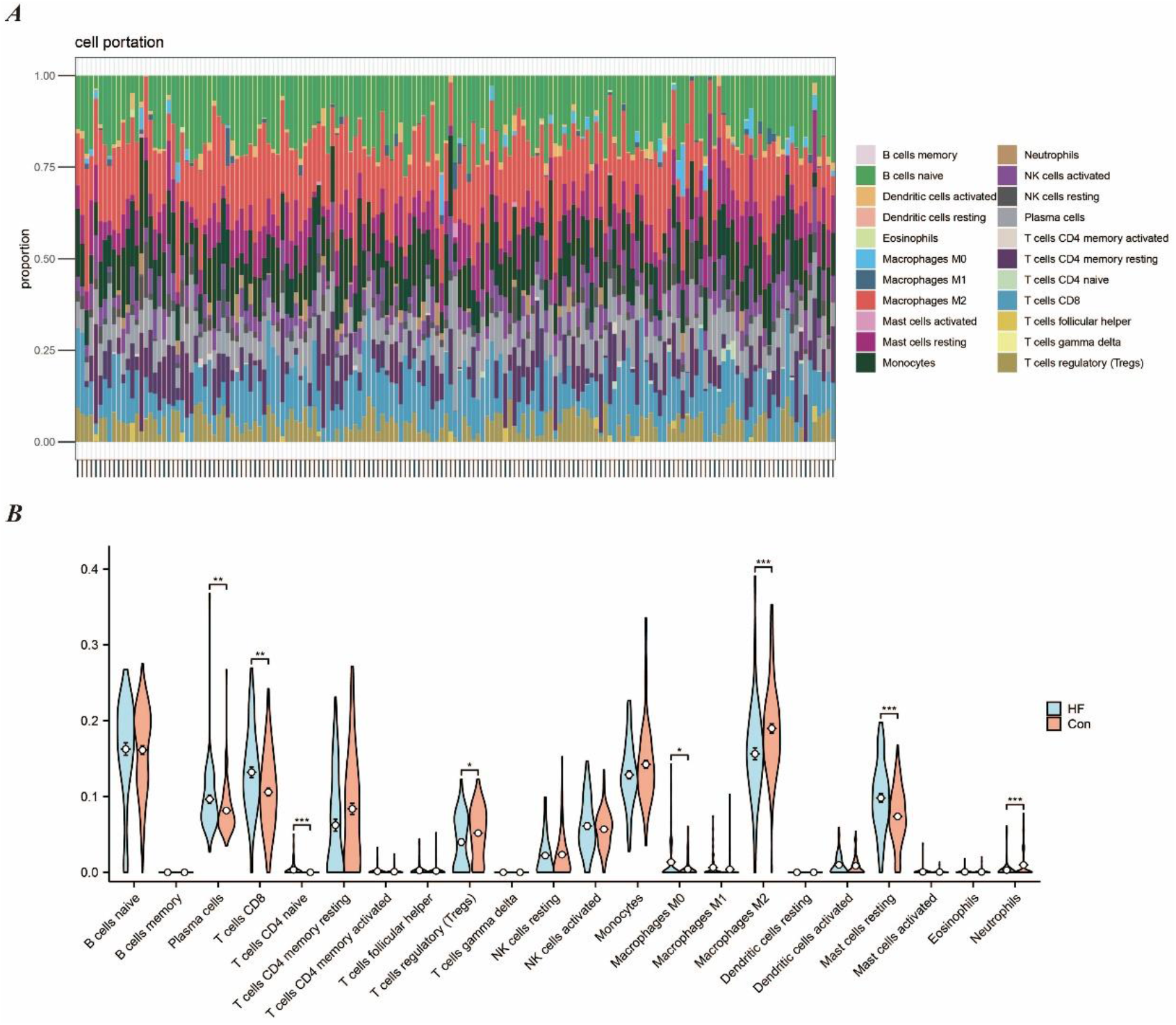
Immune cell infiltration analysis between HF and control in GSE57338-train. (A) The proportion of 22 kinds of immune cells in different samples visualized from the bar plot. (B) Expression of the 8 dysregulated immune cells in the HF and controls as seen in the violin plot.

## DISCUSSION

Winter is a peak season for cardiovascular diseases, and infectious diseases such as influenza also tend to be prevalent during this time. Therefore, the importance of special care for heart failure patients in winter cannot be overstated. Respiratory infectious diseases such as COVID-19, influenza, and CAP have high incidence and mortality rates, potentially leading to severe complications that exacerbate the overall health burden. For instance, CAP is associated with a range of cardiac complications, including arrhythmias, heart failure, and acute myocardial infarction, which can result in hospitalization and long-term mortality [29]. Similarly, COVID-19 has been shown to cause severe acute respiratory infections (SARI), with outcomes comparable to other causes of SARI, necessitating prolonged hospital stays and intensive care [30]. Given the significant impact of these respiratory infectious diseases on cardiovascular health, understanding their molecular mechanisms and identifying potential therapeutic targets is crucial.

In this study, we focused on the common molecular characteristics of respiratory infectious diseases and their potential impact on heart failure. By integrating datasets from three major respiratory infectious diseases, we identified 51 specific genes associated with respiratory infections. Enrichment analysis revealed that the shared molecular features of these three respiratory infectious diseases primarily involve innate immune responses, inflammation, and coagulation pathways. Defense responses to various bacteria, innate immune responses in mucosa, and the formation of neutrophil extracellular traps (NETs) are crucial defense mechanisms in respiratory infectious diseases [31]. However, it is important to note that excessive NETs formation can also lead to tissue damage [32]. NOD-like receptors are intracellular pattern recognition receptors that can recognize pathogen-associated molecular patterns and damage-associated molecular patterns. Activation of the NOD-like receptor signaling pathway can trigger inflammatory responses and cell death to combat pathogen invasion [33]. Heparin-binding proteins and serine-type peptidases play roles in regulating inflammation and coagulation. In respiratory infectious diseases, altered activity of these factors may affect the extent of the inflammatory response and tissue repair processes [34].

To understand the mechanisms by which respiratory infectious diseases exacerbate heart failure, we conducted enrichment analysis on 10 key genes. The results indicated that these genes are primarily involved in immune responses following viral infection, cell death, and inflammatory responses. Viral infections activate the body’s immune system to eliminate invading viruses. However, this immune response, while clearing the virus, can also cause damage to the heart. Accumulation of viral antigens and inflammatory cells can directly harm myocardial cells, leading to myocarditis and myocardial cell dysfunction [35]. Additionally, cytokine storms induced by viral infections, such as the excessive release of tumor necrosis factor-alpha (TNF-α) and interleukin-6 (IL-6), can exacerbate cardiac inflammation and injury [36]. Viral infections can also activate host immune cells to release pro-apoptotic signaling molecules, such as Fas ligand and TRAIL, thereby accelerating myocardial cell death [37]. The inflammatory response triggered by viral infections is another key factor in the exacerbation of heart failure. Viral infections activate the host’s innate immune system, leading to the infiltration of numerous inflammatory cells, such as macrophages and neutrophils, into the heart, releasing inflammatory mediators like interleukins and interferons. These inflammatory mediators not only exacerbate cardiac inflammation but also affect the electrophysiological properties and contractile function of the heart, further worsening the symptoms of heart failure [38].

Using machine learning algorithms, we identified RSAD2 and IFI44L as key genes and validated their high accuracy (AUC > 0.7), further demonstrating their potential as biomarkers for disease progression and therapeutic targets. RSAD2, also known as viperin, is an interferon-induced protein containing a radical S-adenosylmethionine (SAM) domain. It plays a crucial role in the innate immune response against viral infections. Studies have shown that RSAD2 exhibits broad-spectrum antiviral activity by inhibiting the replication of various viruses through different mechanisms [39]. Additionally, RSAD2 is involved in regulating immune responses by promoting dendritic cell maturation via the IRF7-mediated signaling pathway [40]. In the development of heart failure, RSAD2 is one of the key genes associated with mitochondrial dysfunction and immune cell infiltration [41]. Therefore, given its importance in the interaction between respiratory infections and heart failure, RSAD2 has the potential to become a therapeutic target. IFI44L is another interferon-induced gene associated with antiviral responses by inhibiting viral RNA synthesis [42]. During infection, IFI44L promotes macrophage differentiation and the secretion of inflammatory cytokines, thereby exacerbating myocardial injury [43]. Moreover, ssGSEA results suggest that IFI44L may also be related to myocardial contractile function.

Based on the significant roles of RSAD2 and IFI44L in respiratory infectious diseases and heart failure, we used the DSigDB database to predict six potential therapeutic drugs: acetohexamide, Gadodiamide hydrate, suloctidil, 3’-Azido-3’-deoxythymidine, testosterone enanthate, and tamoxifen. Currently, there is no direct evidence indicating the efficacy of the sulfonylurea hypoglycemic agent acetohexamide in treating respiratory infections and heart failure, but controlling blood glucose may indirectly improve the prognosis of heart failure patients [44]. Testosterone enanthate is an androgen used to treat hypogonadism. Studies suggest that testosterone therapy may improve insulin sensitivity and cardiac function in heart failure patients [45]. Tamoxifen, a selective estrogen receptor modulator, has been shown to possess anti-inflammatory properties and may reduce the risk of cardiovascular diseases [46]. Although some drugs show therapeutic potential, most require further research to determine their efficacy and safety.

Our study revealed significant differences in immune cell infiltration between HF samples and healthy controls. Consequently, the impact of respiratory infections on immune cells may exacerbate the progression of HF. Research indicates that myocardial samples from SARS-CoV-2 infection models show a significant increase in T lymphocytes and macrophages, suggesting that SARS-CoV-2 infection induces an excessive inflammatory response, leading to myocardial remodeling and subsequent fibrosis, thereby worsening HF [47]. Additionally, severe COVID-19 patients exhibit dysregulated immune responses, particularly cytokine storms that result in systemic inflammation and multi-organ failure [48]. Dysregulation of monocytes in COVID-19 patients, especially the reduction of the non-classical CD14dimCD16+ subset, is associated with worse clinical outcomes, increasing mortality in patients with respiratory failure and cardiovascular diseases [49]. The immune response in HF patients with respiratory infections becomes more complex due to the dysregulation of regulatory T cells (Treg) and other lymphocyte subsets. Studies have shown that children with congenital heart disease and bronchopneumonia exhibit altered levels of CD3+, CD4+, and CD8+ T cells, indicating impaired cellular immunity, which may predispose them to severe infections and subsequent HF [50]. Furthermore, macrophages have been implicated in cardiac injury during viral ARDS, with an increase in CCR2+ macrophages leading to cardiac inflammation and dysfunction [51]. Therefore, understanding the immune status of HF patients in the context of respiratory infections is crucial. The significant differences in immune cell infiltration and associated inflammatory responses provide deeper insights into the mechanisms by which respiratory infections exacerbate HF, paving the way for the development of targeted therapies aimed at modulating immune responses to improve clinical outcomes in HF patients.

The novelty of our study lies in several key aspects. First, we identified the common molecular characteristics of respiratory infectious diseases and their impact on heart failure using bioinformatics approaches. Subsequently, we pinpointed the key genes exacerbating heart failure due to respiratory infectious diseases through three machine learning algorithms and validated these findings across multiple external datasets. We identified six potential therapeutic drugs using the DSigDB database. Finally, we assessed the impact of immune cells on the myocardium, which aids in understanding the mechanisms by which respiratory infections worsen HF.

Despite these advancements, our study has several limitations. Firstly, it remains unclear whether the elevated mRNA levels will lead to a parallel increase in protein expression, as many biological functions are executed through post-translational modifications. Secondly, although we validated our findings across multiple datasets, further animal experiments and clinical trials are necessary to confirm our results. Lastly, our predicted drugs and immune-targeted therapies have not been validated for relevance and efficacy in clinical settings, necessitating future integration of clinical trials to enhance the reliability of our findings.

## CONCLUSION

Our study successfully identified the common molecular characteristics of respiratory infectious diseases (COVID-19, influenza, and community-acquired pneumonia) and their potential impact on heart failure. Through differential expression analysis, WGCNA, and machine learning algorithms, we pinpointed key genes that may exacerbate heart failure in the context of respiratory infectious diseases. Enrichment analysis and ssGSEA provided insights into the biological processes and pathways involving these genes. Immune infiltration analysis helped us understand the mechanisms by which respiratory infections worsen HF. Finally, we predicted potential therapeutic drug molecules using the DSigDB database. Overall, our findings contribute to a better understanding of the molecular interactions between respiratory infectious diseases and heart failure, paving the way for future research and therapeutic strategies.

## Acknowledgements

Thanks to Figdraw and Servier Medical Art for providing the drawing materials.

## CRediT authorship contribution statement

Yiding Yu: Conceptualization, Methodology, Validation, Formal analysis, Investigation, Resources, Visualization, Writing – original draft, Writing – review & editing.

Juan Zhang: Supervision, Software, Data curation, Writing – review & editing.

Jingle Shi: Supervision, Software, Writing – review & editing.

Huajing Yuan: Methodology, Validation, Writing – review & editing.

Quancheng Han: Investigation, Resources, Writing – review & editing.

Yitao Xue: Project administration, Writing – review & editing.

Yan Li: Project administration, Writing – review & editing, Funding acquisition.

## Consent for publication

Not applicable.

## Ethics approval and consent to participate

Not applicable.

## Data availability statement

Publicly available datasets were analyzed in this study. This data can be found here: GSE57338, GSE5406, GSE157103, GSE196399, GSE164805, GSE185576, and GSE94916.

